# Laurdan and Di-4-ANEPPDHQ probe different properties of the membrane

**DOI:** 10.1101/076752

**Authors:** Mariana Amaro, Francesco Reina, Martin Hof, Christian Eggeling, Erdinc Sezgin

## Abstract

Lipid packing is a crucial feature of cellular membranes. Quantitative analysis of membrane lipid packing can be achieved using polarity sensitive probes whose emission spectrum depends on the lipid packing. However, detailed insight into the exact mechanism that causes the spectral shift is essential to interpret the data correctly. Here, we analysed frequently used polarity sensitive probes, Laurdan and di-4-ANEPPDHQ, to test whether the underlying physical mechanisms of their spectral shift is the same, thus whether they report on the same physico-chemical properties of the cell membrane. Their steady-state spectra as well as time-resolved emission spectra in solvents and model membranes showed that they probe different properties of the lipid membrane. Our findings are important for the application of these dyes in cell biology.

## Introduction

The cellular membrane is a fluid structure [1] that is mostly ensured by the complex mixtures of phospholipids and sterols. Cholesterol in eukaryotes, for instance, is the main component that modulates the fluidity of the membranes [2]. When forming a bilayer on their own, most of the saturated lipids form “gel-like” membranes that have extremely slow or almost no diffusion [3]. Cholesterol fluidizes these gel membranes, making them “liquid” membranes [4]. Membrane fluidity is one of the main determinants of the molecular mobility in the membrane, therefore, it is crucial for membrane bioactivity [5, 6].

Membrane fluidity is generally measured by electron spin resonance, nuclear magnetic resonance or diffusion of membrane molecules [7]. One indirect, yet easy way to infer the fluidity of membranes is to use polarity sensitive probes whose emission spectra change with the polarity of the environment [8]. Polarity in biomembranes generally represents the hydration level of the bilayer [8]. Saturated lipids form relatively more tightly packed membranes (or more ordered membranes) where there is less space for water molecules [9]. Thus, there are less water molecules in the hydrophobic/hydrophilic interface of this type of membrane compared to membranes composed of unsaturated lipids which form relatively loosely packed membranes (disordered membranes) [9] and have higher water content in their hydrophobic/hydrophilic interface.

Laurdan is one of the most commonly used probes [10] to discern lipid packing in biomembranes. It has an emission maximum of 440 nm and 490 nm in gel and liquid crystalline phase membranes, respectively [11]. In comparison, the more recent probe di-4-ANEPPDHQ [12] has a red-shifted spectra with emission maximum at 560 nm or 610 nm in gel or liquid crystalline phase membranes, respectively [13]. The spectral shift is used to calculate the generalized polarization (GP), which is a relative quantitative measure for lipid packing [14, 15]. Although the GP index is easy to obtain and extremely useful to demonstrate the relative changes in membrane lipid packing, it may overlook several physical-chemical aspects of the membrane. One specific cause of this may be the exact mechanisms behind the spectral shift of these dyes. Thus, before interpretation of the empiric values obtained using these probes, the mechanisms of their spectral shift have to be thoroughly addressed.

Here, we investigated the processes behind the spectral shift of aforementioned probes (laurdan and di-4-ANEPPDHQ), which are generally used for the same purpose of obtaining an empirical value for lipid packing. The GP of both dyes successfully probes the differences in lipid packing in phase-separated cell-derived giant plasma membrane vesicles (GPMVs). However, they react differently to solvents with varying polarity suggesting that there may lay different mechanisms behind their spectral shift. The GP of laurdan is much more sensitive to temperature differences while GP of di-4-ANEPPDHQ is more sensitive to cholesterol content in well-defined liposome systems in the liquid crystalline phase. Investigations of time-dependent fluorescent shift of both dyes in liposomes demonstrated that there is not a simple dipolar relaxation of the di-4¬ANEPPDHQ dye, in contrast to laurdan. Our findings show that laurdan and di-4-ANEPPDHQ react differently with the biomembranes, thus reflect different features of the membrane. This should be seriously considered when these dyes are applied to cell biology questions where multiple processes, such as hydration, mobility, cholesterol content and membrane potential, can affect the spectra of these probes.

## Materials and Methods

### Materials

Laurdan and di-4-ANEPPDHQ were obtained from ThermoFisher. RBL-2H3 cells were cultured in MEM supplied with 10% FCS and 1% L-glutamine (all supplied from Sigma-Aldrich). Lipids were bought from Avanti Polar Lipids.

### Preparation of giant plasma membrane vesicles

GPMVs were prepared as previously described [16]. Briefly, cells were incubated with 25 mM paraformaldehyde and 10 mM DTT (Sigma) at 37 ºC for 1 h. They were collected and incubated with 250 nM laurdan or di-4-ANEPPDHQ for 30 min at room temperature and then transferred to BSA-coated 8-well glass bottom Ibidi chambers.

### Spectral imaging of vesicles

Laurdan or di-4-ANEPPDHQ labeled GPMVs were imaged using the spectral imaging mode of a Zeiss 780 microscope equipped with a 40x, 1.2NA objective as reported previously in detail [17].

### Spectrophotometry measurements of vesicles

Spectra of laurdan and di-4-ANEPPDHQ in different solvents were measured in a quartz 96-well plate (Hellma). 385 nm and 488 nm were used for excitation of laurdan and di-4-ANEPPDHQ, respectively. Emission was collected between 400-700 nm for laurdan and 500-700 nm for di-4-ANEPPDHQ.

### Preparation of liposomes

Chloroform solutions of POPC (1-palmitoyl-2-oleoyl-*sn*-glycero-3-phosphocholine) and cholesterol were combined in the appropriate amounts. The organic solvents were then evaporated under a stream of nitrogen. For thorough removal of the solvent, the lipid films were left under vacuum overnight. HEPES buffer (10 mM HEPES, 150 mM NaCl, pH 7.0, 0.2 mM EDTA) was then added to the dried lipid film (lipid concentration of 1 mM), which was left to hydrate for 30 minutes. The resulting suspension was vortexed for at least 4 min and then extruded through polycarbonate membranes with a nominal pore diameter of 100 nm (Avestin, Ottawa, Canada). Laurdan (methanol solution) was added to chloroform/lipids solution before the evaporation under nitrogen. Di-4-ANEPPDHQ (aqueous solution) was added at the end of the hydration stage. The molar ratio of fluorescent probes to lipids was 1:100. For measurements, the vesicle suspensions were diluted to an overall lipid concentration of 0.5 mM.

### Liposome fluorescence measurements

The temperature in the cuvette holders was maintained using a water-circulating bath. Steady-state excitation and emission spectra were acquired using a Fluorolog-3 spectrofluorometer (model FL3-11; Jobin Yvon Inc., Edison, NJ) equipped with a xenon arc lamp. The steady-state spectra were recorded in steps of 1 nm (bandwidths of 1.2 nm were chosen for both the excitation and emission monochromators) in triplicate and averaged. Fluorescence decays were recorded on a 5000 U single-photon counting setup using a cooled Hamamatsu R3809U-50 microchannel plate photomultiplier (IBH, Glasgow, U.K.) and either a NanoLED 11 laser diode (375 nm peak wavelength, 1 MHz repetition rate) or a PicoQuant pulsed diode laser (470 nm peak wavelength, 2.5 MHz maximum repetition rate), for measurements with laurdan and di-4-ANEPPDHQ respectively. A 399nm, or 499 nm, cut-off filter was used to eliminate scattered light, for measurements with laurdan and di-4-ANEPPDHQ respectively. The signal was kept below 1% of the repetition rate of the light source. Data were collected until the peak value reached 5000 counts. The full width at half maximum (FWHM) of the instrument response function was 78 ps and 84 ps, for measurements with laurdan and di-4-ANEPPDHQ respectively.

### Time-resolved emission spectra (TRES) measurements

Fluorescence emission decays were recorded at a series of wavelengths spanning the steady-state emission spectrum ([400–550 nm] in steps of 10 nm for Laurdan and [540-700] nm in steps of 14 nm for di-4-ANEPPDHQ). The fluorescence decays were fitted to a multi-exponential function via the deconvolution method using the IBH DAS6 software. Three exponential components were necessary to obtain a satisfactory fit of the data. The purpose of the fit is to deconvolve the instrumental response from the data, which should not be over-parameterized. The fitted decays together with the steady-state emission spectrum were used for the reconstruction of time-resolved emission spectra (TRES) by a spectral reconstruction method [18]. The reconstruction routine was implemented in Matlab.

### GP of laurdan and di-4-ANEPPDHQ

The generalized polarization spectra of Laurdan (GPlaurdan) was calculated according to ref [15]: 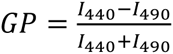 where, *I_440_* and *I_490_* represent the fluorescence intensity emitted at 440 nm and 490 nm, respectively. The generalized polarization spectra of di-4-ANEPPDHQ (GPdi-4) was calculated according to refs [13, 19]: 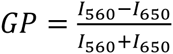, where *I_560_* and *I_650_* represent the fluorescence intensity emitted at 560 nm and 650 nm, respectively.

## Results and Discussion

The cellular plasma membrane is a complex structure inheriting a multitude of different lipids and proteins. Consequently, the spectral emission characteristics of the polarity-sensitive dyes laurdan and di-4-ANEPPDHQ are influenced by several environmental factors such as acyl chain saturation, charge of the head groups, presence of cholesterol, hydration of the membrane and presence of proteins [13, 20–23] and we need reliable knowledge on how these dyes sense their environment, which can best investigated in well-defined systems. We applied fluorescence microscopy and spectroscopy on laurdan and di-4-ANEPPDHQ in solvents and model membranes to investigate their ability to discern lipid packing and to reveal mechanisms behind their environment sensitivity.

### Ability to empirically discern ordered and disordered phases

We first questioned how well both probes empirically distinguish differences in membrane ordered environments in phase-separated model membranes, specifically in cell-derived giant plasma membrane vesicles (GPMVs). We performed spectral imaging where fluorescence images of the equatorial plane of the vesicles were recorded in ≈10 nm wide wavelength intervals, i.e. a fluorescence emission spectrum was acquired for each image pixel with 10 nm spectral resolution (Figure 1a) and thus separate spectra in the ordered and disordered phases were obtained (Figure 1b, c). Laurdan displayed a higher intensity in the liquid ordered phases at all wavelengths, while that of di-4-ANEPPDHQ was stronger in the liquid disordered phases (Figure 1a). Both dyes exhibited a significant spectral shift between the ordered and disordered environment (Figure 1b, c) from which the generalized polarization index (GPlaurdan and GPdi-4) was calculated (Figure 1d). The final GP map (i.e. the GP values determined for each image pixel) of the vesicles showed an efficient distinction between ordered and disordered phases for both probes. We conclude that both probes are empirically capable of distinguishing ordered vs disordered membrane environments in model membranes.

**Figure 1.**
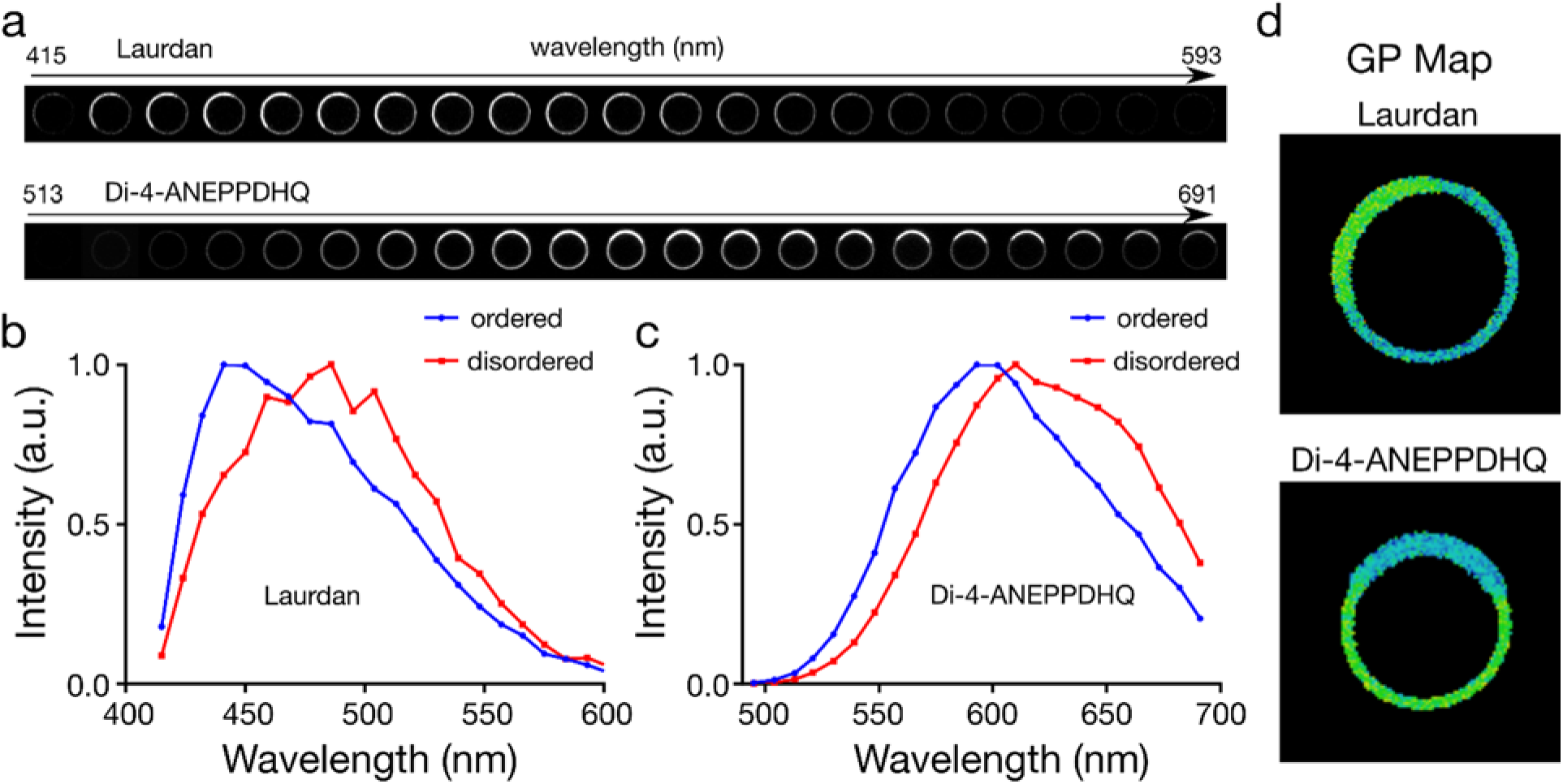
GP imaging of laurdan and di-4-ANEPPDHQ in phase separated GPMVs. a) Representative spectral images, b, c) fluorescence emission spectra in b) ordered and c) disordered domains of GPMVs, and d) GP maps of a phase separated GPMV doped with laurdan or di-4-ANEPPDHQ.

### Dependence on solvent polarity

In order to investigate the influence of purely polarity on the emission characteristics of the dyes, we characterized their fluorescence emission spectra in solvents with varying polarity. Unfortunately, there is no universal polarity index for solvents. As an example, we selected three solvents with significantly different polarity, namely ethanol, DMSO and chloroform. These solvents have different polarity values according to different polarity indices (see Table 1), namely the dielectric constant and the Dimroth and Reichard polarity scales. Chloroform is considered the most non-polar solvent in both scales. However, the relative position of DMSO and ethanol depends on which index is considered. DMSO is more polar than ethanol in terms of dielectric constant, while it is less polar than ethanol in the Dimroth and Reichard polarity index (Table 1).

**Table 1.** Polarity indexes of ethanol, DMSO and chloroform according to their dielectric constants and the Dimroth and Reichard polarity scale.

| Solvent | Dielectric constant | Dimroth and Reichardt |
| --- | --- | --- |
| Water (reference) | 78.54 | 63.1 |
| DMSO | 47 | 45.1 |
| Ethanol | 24.6 | 51.9 |
| Chloroform | 4.81 | 39.1 |

The reason for this difference lies in the definition of these polarity scales. The Dimroth and Reichard polarity index is based on the measure of the ionizing power of a solvent; those with ability to form hydrogen bonds have higher solvation capacity, i.e. stabilizing interactions between the solute and the solvent. On the other hand, the dielectric constant is a relative measure of the chemical polarity of a solvent, or its polarizability. Therefore, the polarity scaling according to the dielectric constant does not consider the possibility of hydrogen bonding between solvent and solute.

The spectra of laurdan and di-4-ANEPPDHQ in the three different solvents exhibit different solvatochromic behaviour. While the spectrum of Laurdan was red-shifted in the order λEthanol˃ λDMSO ˃ λChloroform, the spectrum of di-4-ANEPPDHQ followed the order λDMSO˃ λEth˃ λChloroform (Figure 2). This shows that the photophysical behaviour of the two probes is not identical. The characteristics of laurdan follow that expected for a dye obeying the Dimroth and Reichardt index, while that of di-4-ANEPPDHQ is in line with the polarity scale based on the dielectric constants. This different behaviour reflects the ability of laurdan to form hydrogen bonds with the solvent, which does not seem to occur with di-4-ANEPPDHQ. Thus, the solvatochromic behaviour of laurdan follows the scale that accounts for the hydrogen bonding effect, and the solvatochromic behaviour of di-4-ANEPPDHQ correlates simply with the dielectric constants of the solvents. This differential behaviour of Laurdan and di-4-ANEPPDHQ points out that these probes may react differently with the membrane environments.

**Figure 2.**
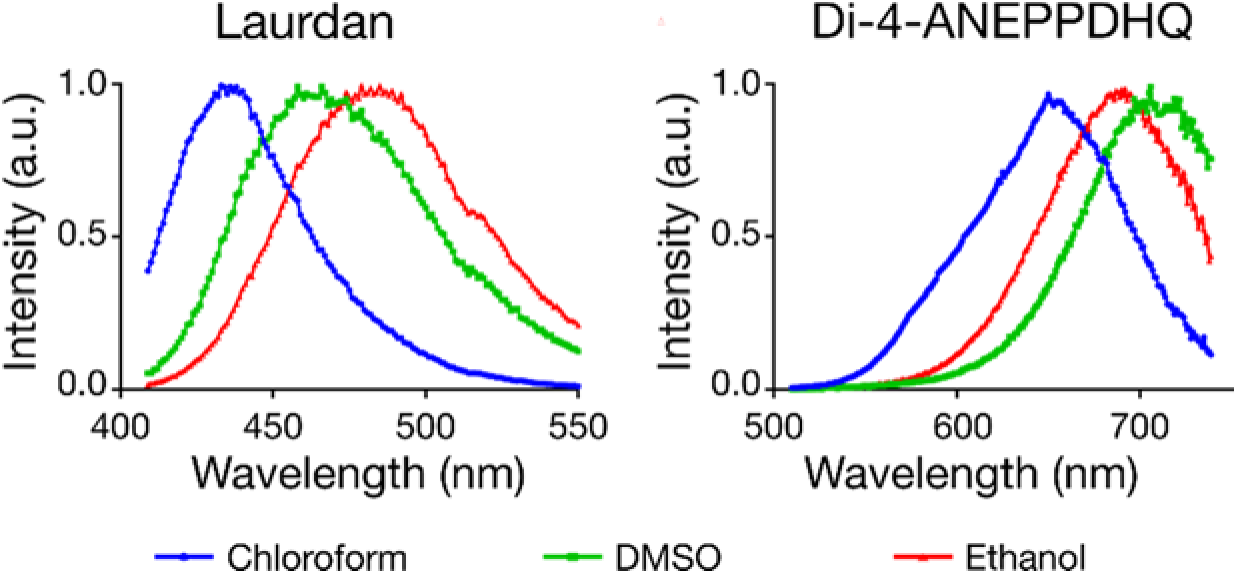
Fluorescence emission spectra of laurdan and di-4-ANEPPDHQ in chloroform (blue), DMSO (green) and ethanol (red).

### Generalized polarisation (GP) in liposomes

Another well-defined system, large unilamellar vesicles (LUVs), was used to investigate how the spectra of laurdan and di-4-ANEPPDHQ dyes are dependent on membrane lipid packing. For this, we acquired fluorescence emission spectra of these dyes in LUVs composed of POPC or POPC + 10% cholesterol (POPC/Chol) at 23 and 37 ºC. From these spectra we determined the generalized polarisation (GP) values of the dyes, GPlaurdan and GPdi-4 (Table 2), the more negative the GP the more disordered or less packed the bilayer (see experimental procedures). The increase in membrane fluidity caused by the increase in temperature is clearly measurable by laurdan, whose GPlaurdan value is a very sensitive indicator in both POPC and POPC/Chol bilayers. In contrast, the GPdi-4 values of di-4-ANEPPDHQ hardly increase with temperature in both POPC and POPC/Chol LUVs. Consequently, GPdi-4 seems to be a less sensitive indicator of changes in membrane order than GPlaurdan. For example, GPlaurdan values in the POPC/Chol bilayer change from 0.026 at 23 ºC to -0.254 at 37 ºC, i.e. ΔGPlaurdan = 0.280, while the GPdi-4 values vary by only ΔGPdi-4 = 0.045. This is a surprising observation, since the lipid packing of the POPC/Chol bilayer is known to be very different between 23 and 37 ºC; it is highly disordered at 37 ºC, while 23 ºC is very close to the transition temperature below which separation into liquid ordered and disordered phases occurs [24].

**Table 2.** GP values of laurdan and di-4-ANEPPDHQ in the different LUV systems. Errors are the standard error of mean.

| Sample<br>Temperature | $GP_{laurdan}$ | | $GP_{di-4}$ | |
| --- | --- | --- | --- | --- |
|  | POPC | POPC/Chol | POPC | POPC/Chol |
| 23 °C | $-0.177 \pm 0.004$ | $0.026 \pm 0.001$ | $-0.346 \pm 0.008$ | $-0.177 \pm 0.008$ |
| 37 °C | $-0.360 \pm 0.003$ | $-0.254 \pm 0.002$ | $-0.397 \pm 0.008$ | $-0.222 \pm 0.006$ |

The observations at different temperatures clearly indicates that the steady-state emission spectrum of di-4-ANEPPDHQ does not report efficiently on the bilayer’s fluidity or lipid packing. However, GPdi-4 is very sensitive on cholesterol, increasing by ΔGPdi-4 = 0.187 (at 23 ºC) and 0.175 (at 37 ºC), even larger than ΔGPlaurdan = 0.151 (at 23 ºC) and 0.106 (at 37 ºC). The sensitivity on cholesterol but insensitivity on temperature suggests that the change in GPdi-4 is caused by a specific effect of cholesterol [23], rather than a consequence of the known cholesterol-induced increase in lipid packing [25–29].

### Time-dependent Fluorescence Shift (TDFS)

In order to investigate in detail the dipolar relaxation properties of di-4-ANEPPDHQ and laurdan, we performed Time-dependent Fluorescence Shift (TDFS) measurements of the dyes in the same LUV systems. It is well known that values of GPlaurdan are affected by both the lipid packing and the hydration level of the membrane bilayer. Nonetheless, TDFS measurements have shown that GPlaurdan reflects predominantly the mobility of the hydrated *sn-*1 carbonyls (i.e. lipid order) and not the extent of hydration of a lipid bilayer in the liquid crystalline phase [20]. Yet, such correlation has not been investigated for di-4-ANEPPDHQ.

TDFS experiments are based on the ultrafast change in the dipole moment of a fluorophore upon electronic excitation, to which its solvation shell must respond. This dipolar relaxation causes a time(*t*)-dependent shift of the peak maximum ν(t) of the emission spectrum, i.e. TDFS probes the time-resolved emission spectra (TRES). The analysis of ν(t) uniquely reveals independent information on the polarity (hydration) and on the molecular mobility in the immediate environment of the dye. The total amount of fluorescence shift Δν is proportional to polarity [18], and is calculated as:

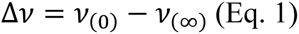

where *v*(0) is the position of TRES maximum at t=0 estimated using the method of Fee and Maroncelli [30] and *v*(∞) is the position of the TRES at the fully relaxed state. The TDFS kinetics depend on the dynamics [31] of the polar moieties in the vicinity of the probe and can be expressed as the integrated relaxation time 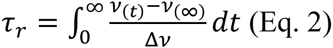

In the particular case of di-4-ANEPPDHQ, we were unable to estimate *v*(0) since this dye is insoluble in non-polar solvents. Therefore, it is not possible to calculate the total amount of fluorescence shift, Δ*v*, or the integrated relaxation time, τr. The latter problem can be circumvented by inspecting the spectral half widths (full width at half-maximum, FWHM) of the reconstructed TRES. The evolution of the FWHM values with time provides useful information on the observed dipolar relaxation phenomenon. It has been shown that the spectral half width, i.e. the FWHM value passes through a maximum during the solvation process [32–34], which is in line with a non¬uniform distribution of solvent response times [34, 35]. The solvent shells of individual fluorophores distributed in a spatially heterogeneous system respond to changes in the local electric fields at different rates. This creates a transient distribution of phases of the individual dipolar relaxation phenomena. The heterogeneity increases significantly during the relaxation process and then decreases once the fluorophores reach the equilibrated excited state. The FWHM of the TRES gives, therefore, a measure of the heterogeneity of the microenvironment of the dye and shows if the entire dipolar relaxation phenomenon was captured within the time-frame of the measurement [18, 32, 36]. Moreover, the time at which the FWHM time-evolution reaches its maximum is a good estimate of the average time taken to complete the dipolar relaxation process and therefore can be used in place of the integrated relaxation time, τr.

### TDFS of laurdan

The values of the fluorescence shift Δ*v* did not significantly shift for the laurdan in the POPC and POPC/Chol LUVs at both temperatures (Figure 3a, Table 3). This implies that the polarity probed by laurdan is identical for the two different lipid bilayers, at both temperatures. However, the relative temporal kinetics are significantly different, e.g. revealed by the temporal development of the maximum or the spectral half width FWHM of the TRES (Figure 3b, Table 3). Table 3 lists the time at which the values of the FWHM of the TRES of laurdan reach their maximum. These values correlate well with the values of GPlaurdan (compare Table 2). It becomes obvious that: **a)** the average time for the completion of the relaxation process is shorter at the higher (37 ºC) temperatures (compared to 23 ºC), i.e. there is a higher mobility of the polar moieties in the vicinity of the probe and the lipid packing is decreased at higher temperature; **b)** addition of cholesterol slows down the process of dipolar relaxation, i.e. there is an increased rigidity in the vicinity of the probe and higher packing of lipids in the cholesterol containing bilayers. These data show that, while TDFS measurements are more informative than GP values, the values of GPlaurdan is still a very good indicator of the order of a lipid bilayer in the liquid crystalline phase.

**Figure 3.**
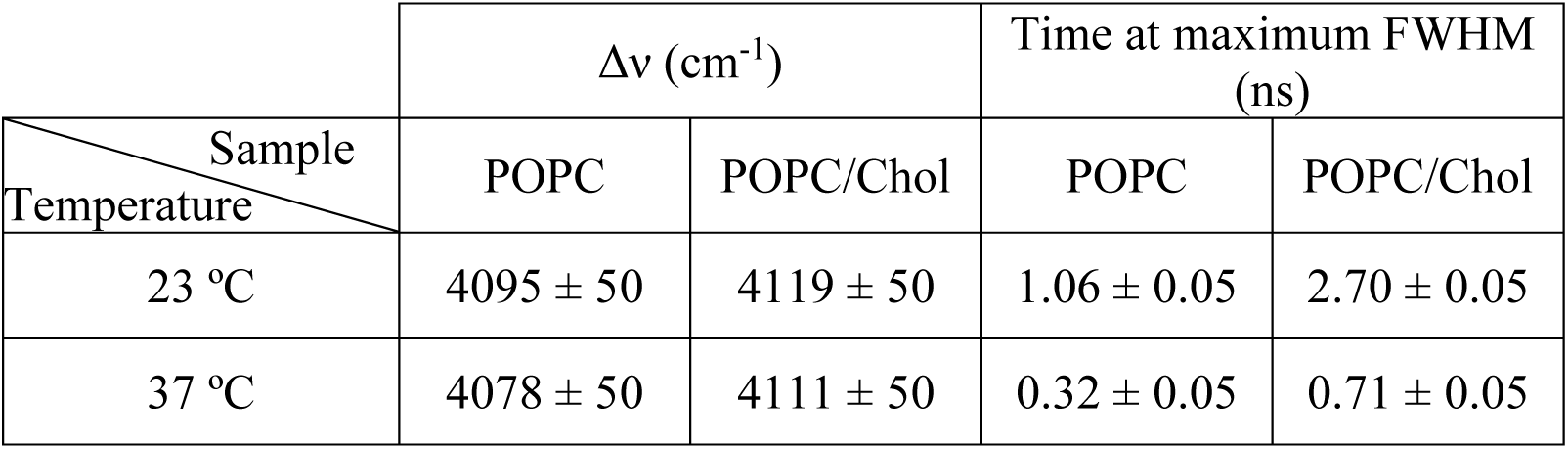
Time-dependence of parameters of the time-resolved emission spectra (TRES) for laurdan in the various liposome samples: a) maximum of TRES, b) full width at half maximum of TRES.

**Table 3.** Values of the fluorescence shift Δ*v* and times for reaching the maximum spectral width FWHM of TRES for laurdan in the different LUV systems. Errors are the intrinsic measurement errors.

| Sample<br>Temperature | $\Delta\nu$ (cm<sup>-1</sup>) | | Time at maximum FWHM<br>(ns) | |
| --- | --- | --- | --- | --- |
|  | POPC | POPC/Chol | POPC | POPC/Chol |
| 23 °C | 4095 ± 50 | 4119 ± 50 | 1.06 ± 0.05 | 2.70 ± 0.05 |
| 37 °C | 4078 ± 50 | 4111 ± 50 | 0.32 ± 0.05 | 0.71 ± 0.05 |

### TDFS of di-4-ANEPPDHQ

Figure 4 shows the time dependence of the TRES maximum for di-4-ANEPPDHQ in the POPC and POPC/Chol bilayers at both 23 ºC and 37 ºC. Due to reasons outlined before, it is not possible to estimate the Δ*v* for this dye. One remedy to this limitation may be to correlate the differences in the energy of the equilibrated excited state, i.e. the asymptote *v*(∞) of the TRES (Figure 4) to the Δ*v*, which would for example indicate that di-4-ANEPPDHQ shows significant differences in hydration with temperature (Table 4). However, the energy *v*(0) of the Frank-Condon state is not the same in all cases. In fact, constant values of *v*(0) in all cases would contradict the results on hydration of laurdan, as revealed by the TDFS experiments. Moreover, from both the solvatochromic behaviour of laurdan and di-4-ANEPPDHQ (Table 1) and from the inability of di-4-ANEPPDHQ to form hydrogen bonds one expects that di-4-ANEPPDHQ has a lower sensitivity to changes in polarity compared to laurdan. Please note that the presence of cholesterol shifts the TRES of di-4-ANEPPDHQ to higher energies, which is consistent with previously reported steady-state fluorescence data of di-4-ANEPPDHQ [23].

**Figure 4.**
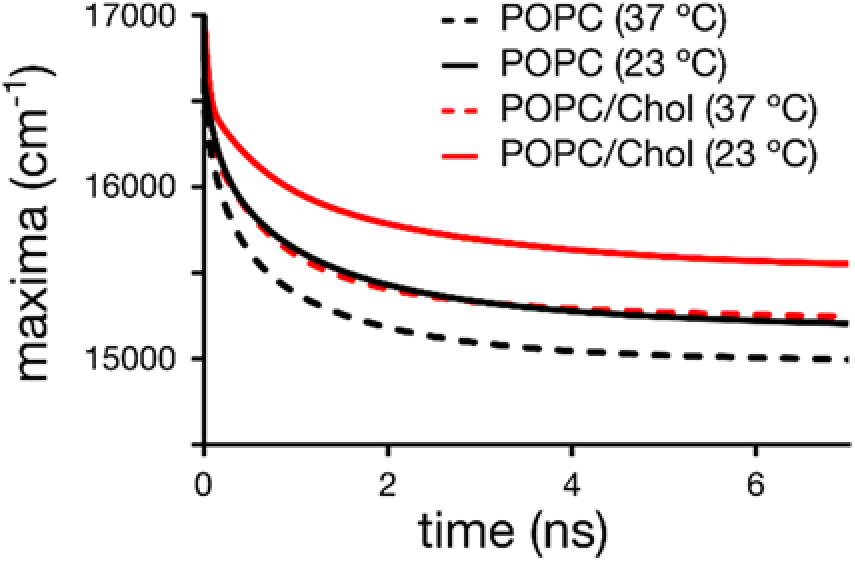
Time-dependence of the maximum of the time-resolved emission spectra (TRES) for di-4-ANEPPDHQ in the different liposome samples.

**Table 4.** Values of *v*(∞) and times for reaching the maximum spectral width FWHM of TRES for di-4-ANEPPDHQ in the different LUV systems. Errors are the intrinsic measurement errors.

| Sample<br>Temperature | $\nu(\infty)$ (cm<sup>-1</sup>) | | Time at maximum FWHM<br>(ns) | |
| --- | --- | --- | --- | --- |
|  | POPC | POPC/Chol | POPC | POPC/Chol |
| 23 °C | 15181 ± 50 | 15533 ± 50 | 1.14 ± 0.05<br>5.22 ± 0.05 | 0.69 ± 0.05<br>4.08 ± 0.05 |
| 37 °C | 14989 ± 50 | 15208 ± 50 | 0.90 ± 0.05<br>4.86 ± 0.05 | 0.68 ± 0.05<br>2.00 ± 0.05 |

The time-evolution of the FWHM values of the TRES for di-4-ANEPPDHQ exhibits complex photophysical characteristics of the dye at all conditions (Figure 5). Instead of the usual single maximum as observed for laurdan, multiple maxima are detected, which suggests the existence of several underlying processes; 1) a very fast process occurs at times <0.1ns which is hardly resolved by our instrument due to missing temporal resolution. Still, our data indicates no significant difference in the fast dynamics between the different LUV samples. Such a fast kinetics is likely to be due to an intramolecular process rather than an effect of the environment of the dye. 2) The presence of two additional maxima at longer times suggests an additional process due to the dipolar relaxation and/or different locations of the dye within the lipid bilayer.

**Figure 5.**
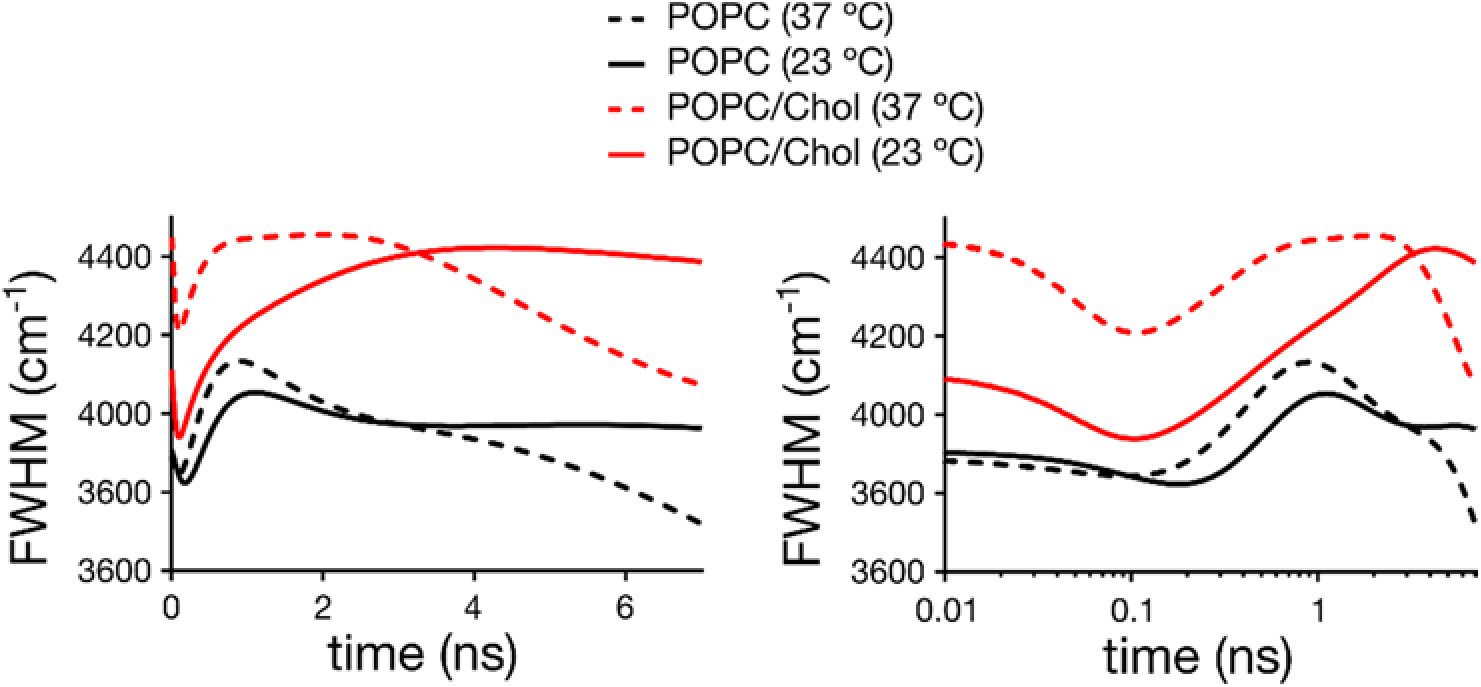
Time-dependence of the values of the spectral width FWHM of the TRES for di-4-ANEPPDHQ in in the different LUVs: (left) linear and (right) logarithmic time scale.

The maxima at the longest time are most sensitive to temperature, shifting for example for POPC/Chol LUVs from 4.08 ± 0.05 ns at 23 ºC to 2.00 ± 0.05 ns at 37 ºC (Table 4). In fact, the shift with temperature is stronger for POPC/Chol compared to POPC LUVs, most probably since the lipid order changes more dramatically for POPC/Chol. In contrast, the peak at around 1 ns hardly changes with temperature in POPC/Chol and POPC LUVs. These observations suggest that the process that causes the slowest modulation of the FHWM (at around 5 ns), likely stems from the dipolar relaxation of the dye. Interestingly, while temperature significantly affects the kinetics of this process, it hardly affects the values of GPdi-4 (compare Table 2). This is contradictory, since changes in the relaxation kinetics do reflect changes in membrane lipid order for laurdan, yet we can conclude that the GPdi-4 parameter is not a good enough indicator of the order of a lipid bilayer in the liquid crystalline phase.

The addition of cholesterol to the lipid bilayers causes an unexpected effect in the kinetics of the relaxation process of di-4-ANEPPDHQ. Instead of slowing down the dynamics of the observed processes (due to the known increase in lipid packing with cholesterol addition), we observed faster relaxation dynamics. The fastest process at < 0.1 ns seems unaffected (at least as far as TRES can resolve, Figure 5). However, the other two maxima are both shifted towards shorter times (Figure 5, Table 4). The faster kinetics may be explained by a different positioning of di-4-ANEPPDHQ in the presence of cholesterol. If, for instance, the dye is pushed slightly out of the membrane upon addition of cholesterol, this would increase the mobility of its immediate environment [37]. Yet, the faster relaxation kinetics (i.e. obviously decreased lipid packing) of di-4-ANEPPDHQ upon cholesterol addition is in contrast with the increase in GPdi-4 values. Most probably, the observed increase in GPdi-4 values is related to a specific influence of cholesterol on the energy levels of the Frank-Condon (*v*(0)) and fully relaxed (*v*(∞)) states of di-4-ANEPPDHQ.

It is worth noting that di-4-ANEPPDHQ belongs to a family of dyes with known electrochromic characteristics [38, 39]. These electrochromic characteristics introduce an enhanced sensitivity to membrane potential, i.e. a change in the voltage or electric field across the membrane will cause a spectral shift in the fluorescence emission, which results from a direct interaction between the electric field and the dipole moments of the ground and excited states. This electrochromic shift is present in both the absorption and emission spectra [40]. Cholesterol is known to either directly or indirectly (via its induced change in lipid packing) increase the internal electrical dipole potential of a lipid membrane bilayer by changing the density of dipoles and the dielectric constant of the membrane interface region [41–43].

## Conclusion

Structural heterogeneity is crucial for the functionality of the cell membrane. It is, therefore, necessary to develop probes and techniques to elucidate the nature of this heterogeneity. The most prominent manifestation of this heterogeneity are spatial differences in the lipid order or packing such as lipid driven separation into ordered and disordered phases as observed in model membranes, which can quantitatively be studied using polarity sensitive probes such as laurdan and di-4-ANEPPDHQ. These dyes change their fluorescence emission spectrum depending on the ordering of the lipid environment. However, it is not clear if they physically probe the same molecular phenomena in the cell membrane. Here, we investigated 1) their ability to discern lipid packing in model membranes, and 2) the photophysical mechanisms behind the observed spectral shifts. Our measurements reveal that both dyes exhibit a sufficiently large spectral shift for discerning ordered and disordered membrane environments in phase separated membranes. Yet, laurdan displays a significantly higher sensitivity for sensing differences in the packing of lipid bilayers in the liquid crystalline phase. TDFS measurements showed that the resulting GPlaurdan values correlate well with the time-scales of the dipolar relaxation processes, which are known to be dependent on the lipid packing of the membrane. Thus, GPlaurdan is an accurate and sensitive indicator of lipid order. On the other hand, the results for di-4-ANEPPDHQ dye revealed complex relaxation kinetics involving multiple processes. The GPdi-4 values do not correlate with lipid packing and are influenced by cholesterol in a specific way. It is of particular importance that di-4-ANEPPDHQ is an electrochromic dye, i.e. its fluorescence emission spectrum is sensitive to the membrane potential. For example, the transmembrane potential of the plasma membrane range from around –40 mV to –80 mV, while that of the mitochondrial membrane is around -140 to -180 mV. Consequently, the electrochromic property of di-4-ANEPPDHQ can substantially bias the interpretation of empiric values when applied to cell biology questions. Therefore, GPdi-4 seems not to be the best indicator of lipid membrane order.

## Acknowledgement

We thank Dr. P. Jurkiewicz for helpful discussions on the TDFS results. This work was supported by the Wolfson Foundation, the Medical Research Council (MRC) (Grant MC_UU_12010/Unit Programmes G0902418 and MC_UU_12025), MRC/BBSRC/ESPRC (Grant MR/K01577X/1), and the Wellcome Trust (Grant ref 104924/14/Z/14). E.S. was supported by EMBO Long Term (ALTF 636-2013) and Marie Curie Intra-European Fellowships (MEMBRANE DYNAMICS).

MH and MA are supported by Grant Agency of CR (P208/12/G016) and they thanks Czech Academy of Sciences for Praemium Academie award.

